# Parallel and non-parallel changes of the gut microbiota during trophic diversification in repeated young adaptive radiations of sympatric cichlid fish

**DOI:** 10.1101/793760

**Authors:** Andreas Härer, Julián Torres-Dowdall, Sina Rometsch, Elizabeth Yohannes, Gonzalo Machado-Schiaffino, Axel Meyer

**Affiliations:** Department of Biology, University of Konstanz, Germany; Zukunftskolleg, University of Konstanz, Konstanz, Germany; Department of Functional Biology, University of Oviedo, Oviedo, Spain

**Keywords:** *Amphilophus citrinellus*, eDNA, stable isotopes, Neotropical cichlids, rapid adaptation

## Abstract

Recent increases in understanding the ecological and evolutionary roles of microbial communities has underscored their importance for their hosts’ biology. Yet, little is known about gut microbiota dynamics during early stages of ecological diversification and speciation. We studied the gut microbiota of extremely young adaptive radiations of Nicaraguan crater lake cichlid fish (*Amphilophus* cf. *citrinellus*) to test the hypothesis that parallel evolution in trophic ecology is associated with parallel changes of the gut microbiota. Bacterial communities of the water (eDNA) and guts were highly distinct, indicating that the gut microbiota is shaped by host-specific factors. Across individuals of the same crater lake, differentiation in trophic ecology was associated with gut microbiota differentiation, suggesting that diet affects the gut microbiota. However, differences in trophic ecology were much more pronounced across than within species whereas little evidence was found for similar patterns in taxonomic and functional changes of the gut microbiota. Across the two crater lakes, we could not detect evidence for parallel changes of the gut microbiota associated with trophic ecology. Similar cases of non-parallelism have been observed in other recently diverged fish species and might be explained by a lack of clearly differentiated niches during early stages of ecological diversification.

## Introduction

The importance of microorganisms for many aspects of their hosts’ biology is increasingly recognized for a wide range of animals, from insects to mammals (1-3). The gut microbiota is a complex and dynamic community that is fundamental for physiological processes, such as regulation of the immune system (4) and nutrient metabolism (5). Further, the significance of microbes in animal evolution has become increasingly appreciated (1, 6, 7). In some cases, divergence of the gut microbiota appears to be strongly correlated with their host’s phylogeny and genetic divergence (8, 9). These findings are supported by the fact that host genetics, together with environmental effects such as diet, contributes to shaping and maintaining gut microbiota composition (10-12). However, open questions remain on how closely the gut microbiota matches the biology of its host during ecological diversification and speciation or whether the composition of the gut microbiota could even be predicted based on the ecology of its host. These important questions can best be addressed in a setting where evolution repeated itself, i.e., in pairs of species that evolved in parallel. Cases of parallel evolution of host species associated with divergence in trophic ecology and habitat use allow us to ask whether the gut microbiota also changes in a predictable and parallel manner. This question has been addressed in several fish species covering a wide range of divergence times but results have been inconsistent. African cichlids from two old adaptive radiations of Barombi Mbo (0.5 - 1 myr) and Tanganyika (9 - 12 myr) show parallel changes of the gut microbiota associated with host diet (13). Yet, studies on lineages that diverged more recently like whitefish and Trinidadian guppies did not find evidence for parallelism (14, 15).

Here, we asked whether evolutionary divergence in host species’ trophic ecology predicts the composition of the gut microbiota of Nicaraguan Midas cichlids (*Amphilophus* cf. *citrinellus*), which represent very young adaptive radiations that, notably, occurred in parallel and sympatrically. Currently, there are 13 described species of Midas cichlids (16-19) and their distribution results from independent colonization events from two older great lakes (Lakes Managua and Nicaragua) that are approximately 500,000 years old (17, 20) into several crater lakes (calderas of inactive volcanoes; all crater lakes are between 1,000 and 23,000 years old) (21). Crater lake Midas cichlids differ from their source populations of the great lakes in traits such as body shape and visual sensitivity (22-25). The colonization events of crater lakes Apoyo (colonized from Lake Nicaragua) and Xiloá (colonized from Lake Managua) are estimated to have occurred as recently as 1,700 and 1,300 generations ago, respectively (18, 26). Within these two crater lakes, multiple species of Midas cichlids evolved in sympatry during these extremely short time spans, hence, ten species are endemic to Apoyo (six) and Xiloá (four) (18, 27). Notably, one slender-bodied, limnetic species (*A. zaliosus* in Apoyo and *A. sagittae* in Xiloá) independently evolved in each of the two crater lakes. These elongated limnetic species are not found in the great lakes and inhabit the open water zone that is exclusive to the deep crater lakes. Limnetic species differ distinctly in body shape from several deep-bodied benthic species in their respective crater lakes (18, 22, 25) and feed at a higher trophic level (22) (Fig. 1). Previously, we have shown that gut microbiotas differ between a benthic-limnetic species pair from Apoyo (28).

**Fig. 1:**
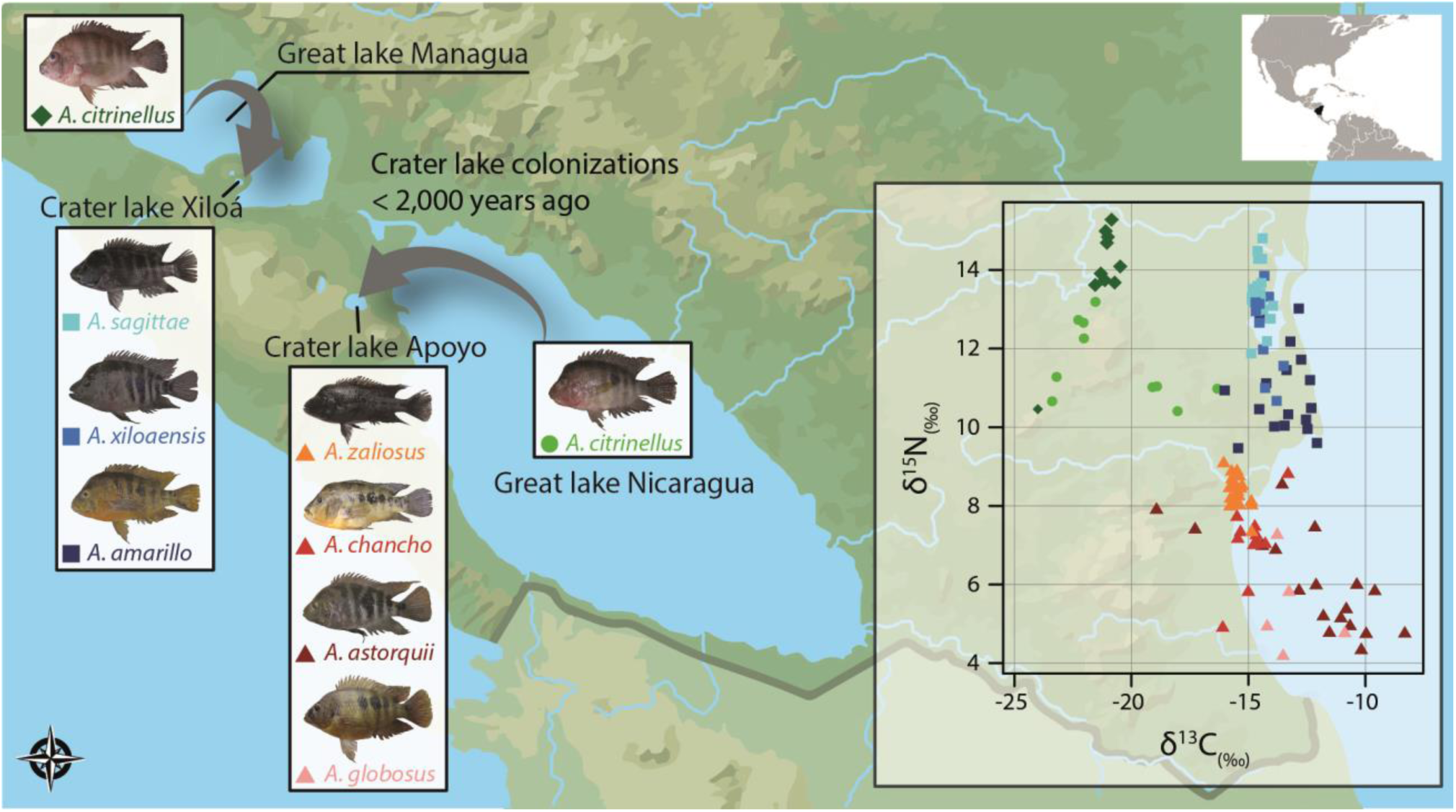
Map of Nicaragua showing the partial distribution of Midas cichlids in Nicaragua. Midas cichlids from the two great lakes (Managua and Nicaragua) colonized crater lakes Apoyo and Xiloá. In these two young crater lakes, multiple endemic species evolved in sympatry, representing a compelling case of parallel adaptive radiations. Within each crater lake, several deep-bodied, benthic and one elongated, limnetic species (*A. sagittae* in Xiloá and *A. zaliosus* in Apoyo) occur, which evolved rapidly within less than 2,000 generations. White inset: Stable isotope analysis of carbon (δ^13^C) and nitrogen (δ^15^N) based on muscle tissue. δ^13^C and δ^15^N values vary significantly not only among environments, but also within each of the crater lakes where substantial variation can be observed among species. Within each of the crater lake, variation in δ^15^N indicates differences in trophic ecology among species.

The extraordinary system of crater lake Midas cichlids is a promising model to elucidate to what extent trophic ecology is mirrored by repeated and parallel changes of the gut microbiota across two very young adaptive radiations. First, we investigated trophic position (stable isotope ratios of carbon and nitrogen) and the gut microbiota of Midas cichlids from the two source lakes, the great lakes Managua and Nicaragua, four species from crater lake Apoyo and three species from crater lake Xiloá. in particular, we tested the hypotheses that (i) species from distinct lakes differ in their gut microbiota but also from the bacterial communities of their natural environments and (ii) repeated adaptation to different trophic niches is associated with parallel changes of the gut microbiota across the two crater lake radiations.

## Methods

### Sample collection

Specimens of the *Amphilophus* cf. *citrinellus* species complex were caught during field trips to Nicaragua in 2014 and 2015 (under MARENA permits DGPN/DB-IC-011-2014 & DGPN/DB-IC-015-2015). We collected *A. citrinellus* populations from the two great lakes, Lake Nicaragua (n = 10) and Lake Managua (n = 10). From two crater lakes, we collected elongated limnetic species, *A. sagittae* from Xiloá (n = 20) and *A. zaliosus* from Apoyo (n = 19), as well as several deep-bodied benthic species, *A. amarillo* (n = 17) and *A. xilaoensis* (n = 16) from Xiloá, *A. astorquii* (n = 19), *A. chancho* (n = 11) and *A. globosus* (n = 5) from Apoyo (Fig. 1). All specimens were sacrificed by applying an overdose of MS-222. Then, whole guts were dissected, cleaned and stored in absolute EtOH at −20°C until DNA extraction. Muscle tissue of the same specimens was collected in absolute EtOH and stored at −20°C for stable isotope analyses. Four technical replicates of environmental DNA (eDNA) were collected along the shores of the four lakes in 2018. Briefly, 500 ml of lake water were filtered through a cellulose nitrate filter (ø 47 mm, pore size 1 µm), stored in Longmire’s solution (29) at −20°C until DNA extraction.

### Trophic analysis

Stable isotope ratios of carbon (δ^13^C) and nitrogen (δ^15^N) were determined based on muscle tissue of the same fish used for gut microbiota analyses. Dried and powdered samples (0.6 mg) were loaded into tin capsules and combusted in a vario Micro cube elemental analyzer (Elementar Analysensysteme, Germany). The resulting gases were fed via gas chromatography into the inlet of a Micromass Isoprime Isotope Ratio Mass Spectrometer (Isoprime, Cheadle Hulme, UK). Two sulfanilamides (Iso-prime internal standards) and two Casein standards were used. Internal laboratory standards indicated measurement errors (SD) of ± 0.03‰ for δ^13^C and 0.12‰ for δ^15^N. Isotopic values are reported in δ-notation in parts per thousand deviations (‰) relative to international standards for carbon (Pee Dee Belemnite, PDB) and nitrogen (atmospheric N2, AIR) according to the following equation:

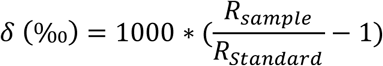

### Library preparation & Illumina Sequencing

DNA from fish guts was extracted from approximately 50-100 mg of tissue from the medial gut section using the commercial QIAamp DNA Stool Mini Kit according to the manufacturer’s protocol (Qiagen, Hilden, Germany). eDNA from water samples was extracted using a QIAGEN DNeasy Blood & Tissue kit. All DNA extractions and PCR amplification were performed under sterile conditions in a laminar flow hood to minimize contamination risk. Concentrations were measured on a Qubit v2.0 Fluorometer (Thermo Fisher Scientific, Waltham, Massachusetts). For each round of extractions, one negative control of sterile H_2_O was included which in no case yielded detectable DNA concentrations. We performed two sequential PCRs and after each one, the amplified product was purified with HighPrep™ PCR beads (MagBio Genomics, Gaithersburg, Maryland). For the first PCR, we used the 515F and 806R primers with a universal 5’ tail, as indicated in the Illumina Nextera library preparation protocol, for DNA amplification of the bacterial V4 region of the 16S ribosomal RNA (292 bp). Briefly, 50 ng (fish guts) or 2 ng (eDNA) of DNA were used as template for the first PCR (2min at 98°C, 10 amplification cycles consisting of 15s at 98°C, 20s at 55°C and 20s at 72°C and a final elongation at 72°C for 2 minutes) and the purified PCR amplicons were the template for the second PCR (2min at 98°C, 20 amplification cycles consisting of 15s at 98°C, 20s at 67°C and 20s at 72°C followed by a final elongation at 72°C for 2 min) using primers including sequencing barcodes as well as the Illumina adapter sequences. Both PCRs were performed in 25 µl reaction volumes, amplifying with the Q5 High-Fidelity polymerase 2x Master Mix (New England Biolabs, Ipswich, MA). After purification, DNA concentrations were measured and specificity of amplification was checked for all samples using gel electrophoresis. Again, a negative control was included during each PCR but no amplified PCR products were detected (based on gel electrophoresis and measured DNA concentrations). Fish gut and eDNA samples were separately pooled in an equimolar manner and size selection was performed on a Pippin Prep device (Sage Science, Beverly, MA). The quality of the pooled libraries was assessed using a Bioanalyzer 2100 (Agilent Technologies, Waldbronn, Germany). Both libraries were paired-end sequenced, each in one lane of the Illumina flow cell. For the fish guts, we sequenced 2×250 bp on an Illumina HiSeq 2500 platform at TUCF Genomics (Tufts University). eDNA was sequenced 2×150 bp on an Illumina HiSeq X-ten at BGI Genomics.

### Gut microbiota analysis

We obtained a total of 62,728,287 (median: 238,073 reads/specimen) and 111,949,556 (median: 6,825,739) raw sequencing reads that could be unambiguously assigned to a specific sample for fish guts and eDNA, respectively. Illumina adapters were removed and reads were trimmed with Trimmomatic v0.36 (30). As there was no overlap between forward and reverse reads for eDNA samples and the sequence quality of forward reads was higher, we used 135 bp of the forward reads for all analyses. The demultiplexed and trimmed reads were imported into the open-source bioinformatics pipeline Quantitative Insights Into Microbial Ecology (QIIME2) (31) to analyze microbial communities of fish guts and water samples. Briefly, sequence quality control was done with the QIIME2 plugin *deblur*. A phylogenetic tree of bacterial taxa was produced with FastTree 2.1.3 (32). Bacterial species richness (total of 3,229 observed OTUs) and bacterial community composition (weighted UniFrac) (33, 34) were calculated and taxonomy was assigned using vsearch (35) against the SILVA 132 ribosomal RNA (rRNA) databases at a 97% similarity threshold (36). The weighted UniFrac distance matrix was visualized with principal coordinate analyses. Since most diversity metrics are sensitive to differences in numbers of reads per sample, we chose 20,000 sequencing reads, the approximate number of reads for the sample with the lowest sequencing depth, as our sampling depth for all further analyses. To determine whether this sampling depth was appropriate to capture a large proportion of the microbial diversity for each sample (bacterial species richness measured as observed OTUs), we rarefied our data (Fig. S1). This analysis confirmed that a large proportion of the gut microbial diversity is already captured at a sequencing depth of 20,000 reads/sample (Fig. S1). Non-parametric Wilcoxon rank-sum tests were used for pairwise comparisons (37) and Kruskal-Wallis tests for comparisons across multiple groups, as implemented in the R stats package (38). To test for gut microbial community differences, both in terms of taxonomic and functional diversity, we applied Permutational Multivariate Analysis of Variance Distance Matrices (PERMANOVA) (39), using the *adonis* function of the R vegan package. Correlations between pairwise distances of stable isotope data (δ^13^C or δ^15^N) and gut microbiotas (weighted UniFrac) among individuals were tested using Pearson’s product-moment correlation. MetaCyc pathway abundances were predicted based on 16S rRNA sequencing data with the PICRUSt2 plugin in QIIME2 (40). Stable isotope ratios were normalized by z-score normalization to test for parallelism across crater lakes. We tested for effects of lake and stable isotope values on OTU and MetaCyc pathway abundance by using linear models (OTU/Metacyc ∼ lake*normalized δ^13^C /δ^15^N). OTU/MetaCyc abundance was scored as parallel across crater lakes if stable isotope values had a significant effect on a given OTU and the interaction term between lake and stable isotope value was non-significant. Only OTUs with a mean proportional abundance of more than 0.1% were selected for the aforementioned analysis. Statistical analyses were performed in R v3.2.3 (41).

## Results

### Diet differentiation among Midas cichlids

In order to obtain information on trophic ecology of all studied Midas cichlid species, we measured stable isotope ratios of carbon (δ^13^C) and nitrogen (δ^15^N). Overall, Midas cichlids from different environments significantly differed in δ^13^C (Kruskal-Wallis test, *P* < 0.001) and δ^15^N (*P* < 0.001; Fig. 1). Within each of the crater lakes, stable isotope ratios differed among species (δ^13^C: *P*_*Apoyo*_ < 0.001, *P*_*Xiloá*_ = 0.001; δ^15^N: *P* < 0.001 for both lakes). These results illustrate that sympatric species of Midas cichlids feed on different carbon sources and further suggests that they also occupy distinctive trophic levels based on nitrogen values (22). As predicted based on diet and inferred trophic niche (22, 27), the limnetic species had the highest nitrogen value in both crater lakes (Fig. 1). Benthic species occupied distinct trophic niches that were generally at lower trophic levels (*A. globosus* in Apoyo, *A. amarillo* in Xiloá) than the limnetic species. Yet, one benthic species largely overlapped with the limnetic species in each crater lake (*A. chancho* with the limnetic *A. zaliosus* in Apoyo, *A. xiloaensis* with the limnetic *A. sagittae* in Xiloá). The benthic *A. astorquii* from Apoyo was highly variable in carbon and nitrogen signatures and largely overlapped with the other species (Fig. 1).

### Gut microbiota differentiation across lakes

Bacterial community composition significantly differed between lake water and fish guts (weighted UniFrac; adonis, *P* = 0.001; Fig. 2), emphasizing that the gut microbiota not merely represents the microbial community of the natural environment. In the water samples, Cyanobacteria (9.9 - 21.9%), Planctomycetes (10 - 23%) and Actinobacteria (9 - 25.1%) constituted a large proportion of microbial communities whereas these groups where much less abundant in the gut microbiota of Midas cichlids (Fig. 3A). In contrast, the gut microbiota was dominated by Proteobacteria (35.4 - 64.9%), Firmicutes (3.9 - 40.4%) and Fusobacteria (2.5 - 21.1%; Fig. 3A), whereas the last two where almost absent in the water. Bacterial species richness (observed OTUs) was significantly higher in water (mean: 1,446 OTUs) compared to fish guts (mean: 448 OTUs) (Wilcoxon rank-sum test, *P* < 0.001; Fig. 3A) and among the water samples, great lakes showed a higher bacterial species richness than crater lakes (*P* < 0.001). Among Midas cichlids, bacterial community composition differed not only across lakes (*P* = 0.001) but also between environment type (great lakes vs. crater lakes; *P* = 0.007). Bacterial species richness was lower in great lake Midas cichlids (*P* < 0.001), however, this pattern disappeared when *A. citrinellus* from Lake Managua was removed from the analysis (*P* = 0.5124). These results clearly show that bacterial species richness is largely constant across Midas cichlids from different environments, except for *A. citrinellus* from Lake Managua that showed strongly reduced bacterial diversity. Next, we investigated whether gut microbiota divergence within crater lakes is associated with the observed differences in stable isotope ratios.

**Fig. 2:**
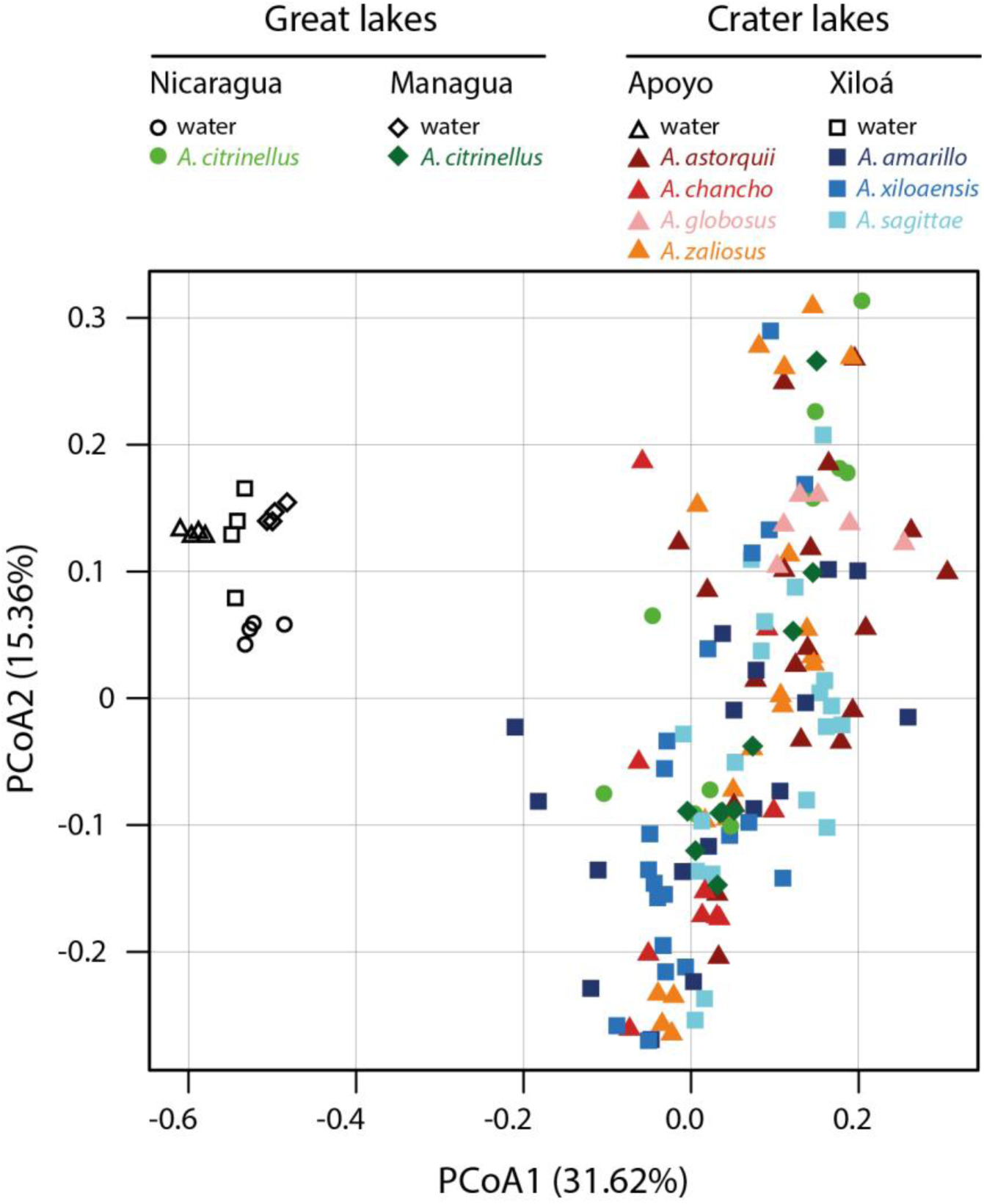
Principal coordinate analysis of bacterial community composition from Midas cichlids’ guts (colored symbols) and their natural environment (eDNA, black symbols) measured as weighted UniFrac. Bacterial communities of eDNA obtained from water samples are clearly differentiated from those of fish guts along PCoA1. Among Midas cichlids, no apparent clustering by lake or species can be detected along PCoAs 1 and 2.

**Fig. 3:**
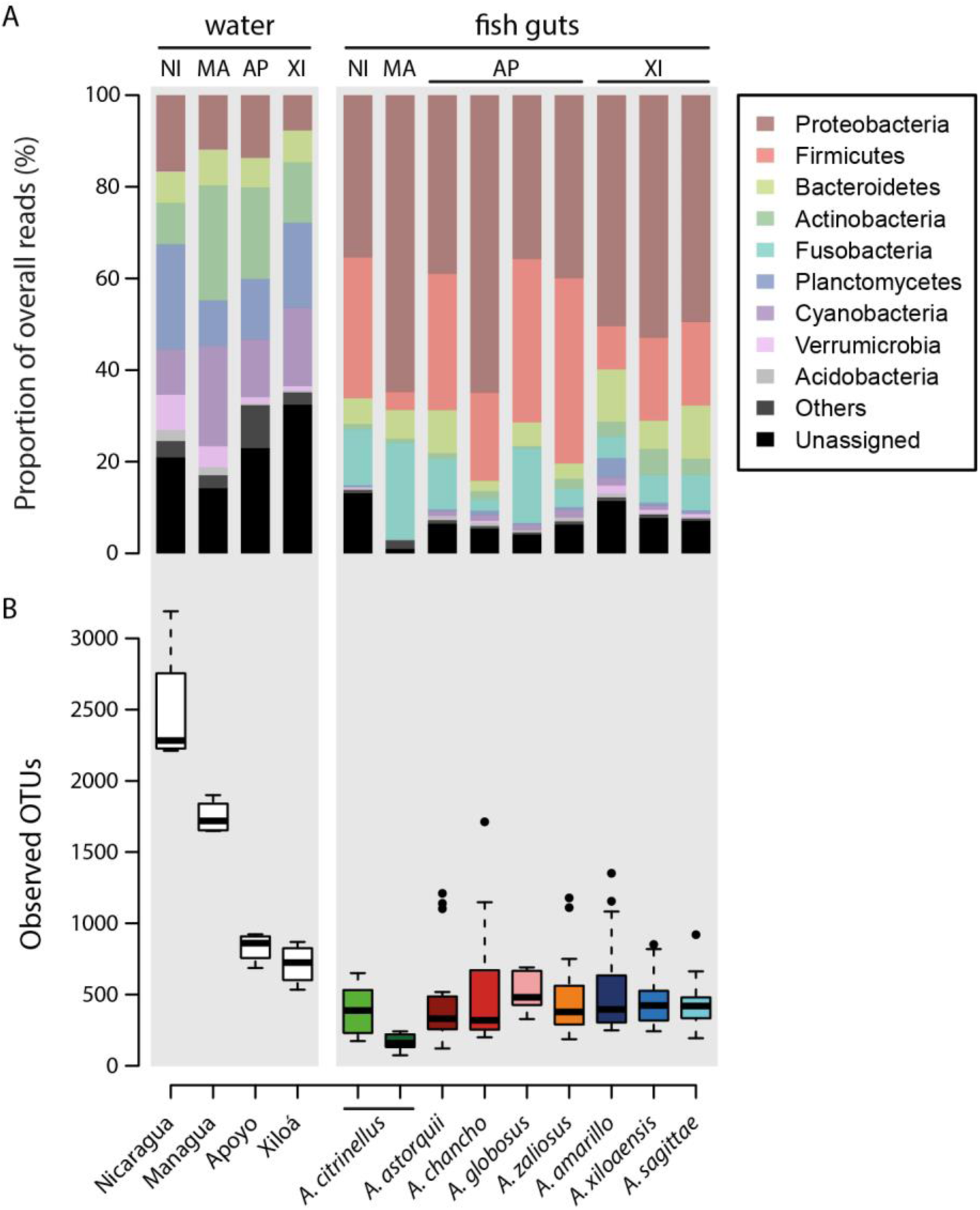
(A) Abundances of the nine most common bacterial phyla found in water and guts of the study species (> 0.5% of overall sequencing reads). The gut microbiota is dominated by Proteobacteria, Firmicutes and Fusobacteria, three phyla which occur only at low abundance in the water. (B) Bacterial species richness (observed OTUs) is higher in the water compared to fish guts. Among Midas cichlid species, there is little variation, only *A. citrinellus* from Lake Managua shows remarkably reduced bacterial species richness.

### Association between trophic ecology and gut microbiota in crater lake adaptive radiations

The two crater lakes are each inhabited by several endemic Midas cichlid species that substantially differ in their morphology, ecological niche, and diet (Fig. 1). Yet, there were no significant differences in bacterial species richness (observed OTUs) among sympatric species within each of the crater lakes (Wilcoxon rank-sum tests, *P* > 0.05 for all pairwise comparisons). Hence, we tested whether bacterial community composition (weighted UniFrac) varied among species of the two parallel crater lake radiations of Apoyo and Xiloá (Fig. S2).

There was overall differentiation in the taxonomic composition of bacterial communities among sympatric species in both crater lakes (*P*_*Apoyo*_ = 0.001, *P*_*Xiloá*_ = 0.007). From the taxonomic composition of the gut microbiota, one can infer the functional bacterial metagenome by predicting the abundance of genes involved in metabolic pathways (40). Predicted functional bacterial metagenomes significantly differed across species only in crater lake Xiloá (*P* = 0.006). Since there is pronounced variation in trophic ecology of both adaptive radiations (Fig. 1), we calculated pairwise distances of trophic ecology (δ^15^N and δ^13^C scores) and correlated these with pairwise distances in bacterial community composition among all individuals within each crater lake (Fig. 4). For carbon, we found a significant positive correlation with bacterial community composition in Apoyo (Pearson’s product-moment correlation, r = 0.092, *P* < 0.001; Fig. 4A) and in Xiloá (r = 0.133, *P* < 0.001, Fig. 4B). For nitrogen, there was a significant positive correlation with bacterial community composition in Xiloá (r = 0.139, *P* < 0.001; Fig. 4D), but only a suggested positive correlation in Apoyo (r = 0.049, *P* = 0.064; Fig. 4C). These results demonstrate that differences in trophic ecology among individuals of the same crater lake are associated with differentiation of the gut microbiota. Distinguishing between intra- and interspecific comparisons of pairwise distances revealed that differentiation in diet is highly distinct among than within species for carbon (Wilcoxon rank-sum test, *P* < 0.001 for both lakes; Fig. 5A) and nitrogen (*P* < 0.001 for both lakes; Fig. 5B). In contrast, taxonomic differentiation of the gut microbiota (*P*_*Apoyo*_ = 0.069, *P*_*Xiloá*_ = 0.004; Fig. 5C) and the predicted functional metagenome (*P*_*Apoyo*_ = 0.824, *P*_*Xiloá*_ = 0.047; Fig. 5D) showed more similar (albeit significantly different in Xiloá) levels between intra- and interspecific comparisons. This clearly illustrates that pronounced differentiation in diet across species is not reflected by equivalent changes of the gut microbiota within crater lakes.

**Fig. 4:**
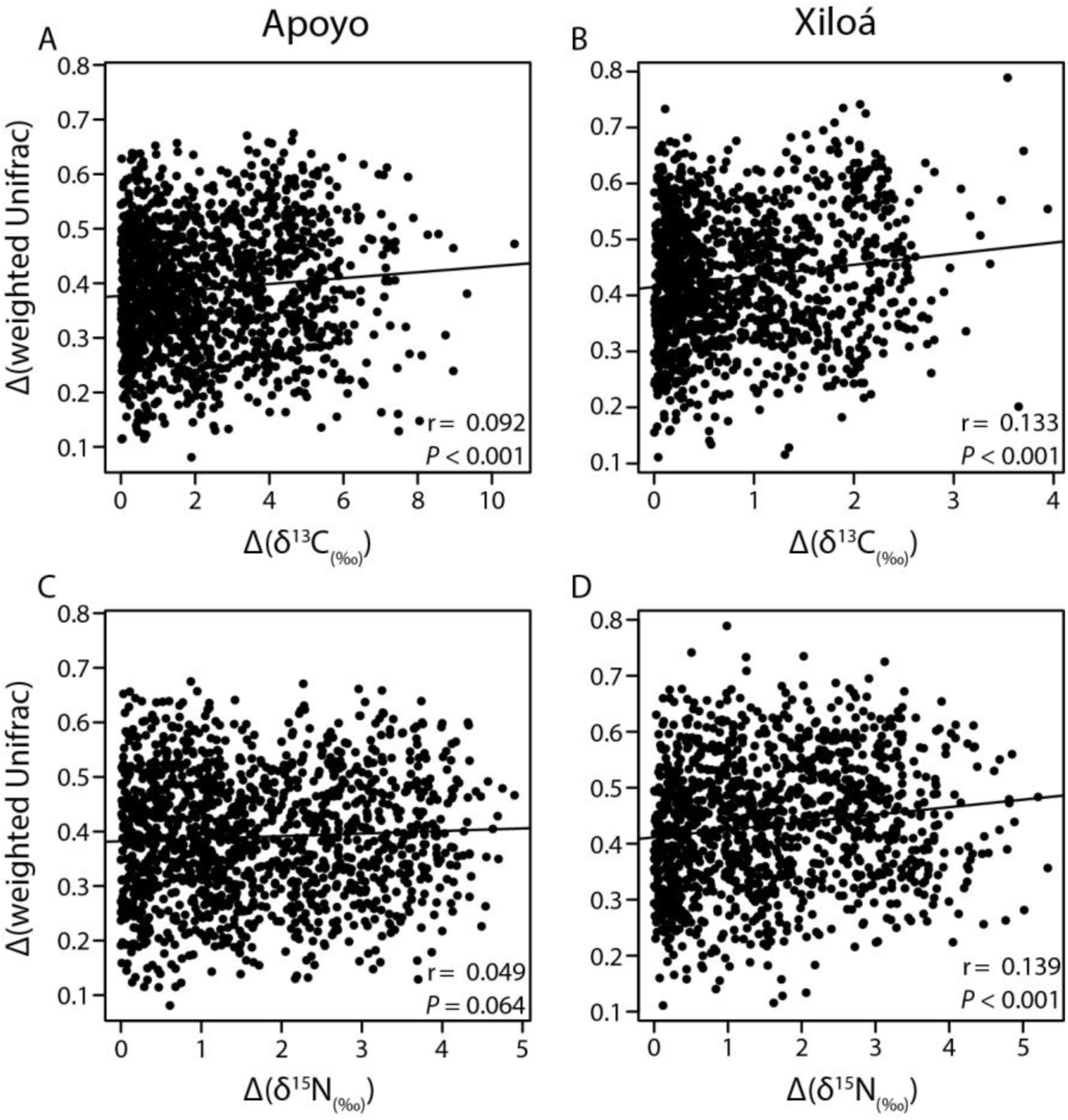
Pairwise distances of gut microbiota composition, Δ(weighted UniFrac), and trophic ecology, Δ(δ^15^N) and Δ(δ^13^C), among all individuals within crater lakes Apoyo (A&C) and Xiloá (B&D). Gut microbiota differentiation is positively correlated with divergence in carbon values in both crater lakes, and with nitrogen values in crater lake Xiloá (Pearson’s product-moment correlation).

**Fig. 5:**
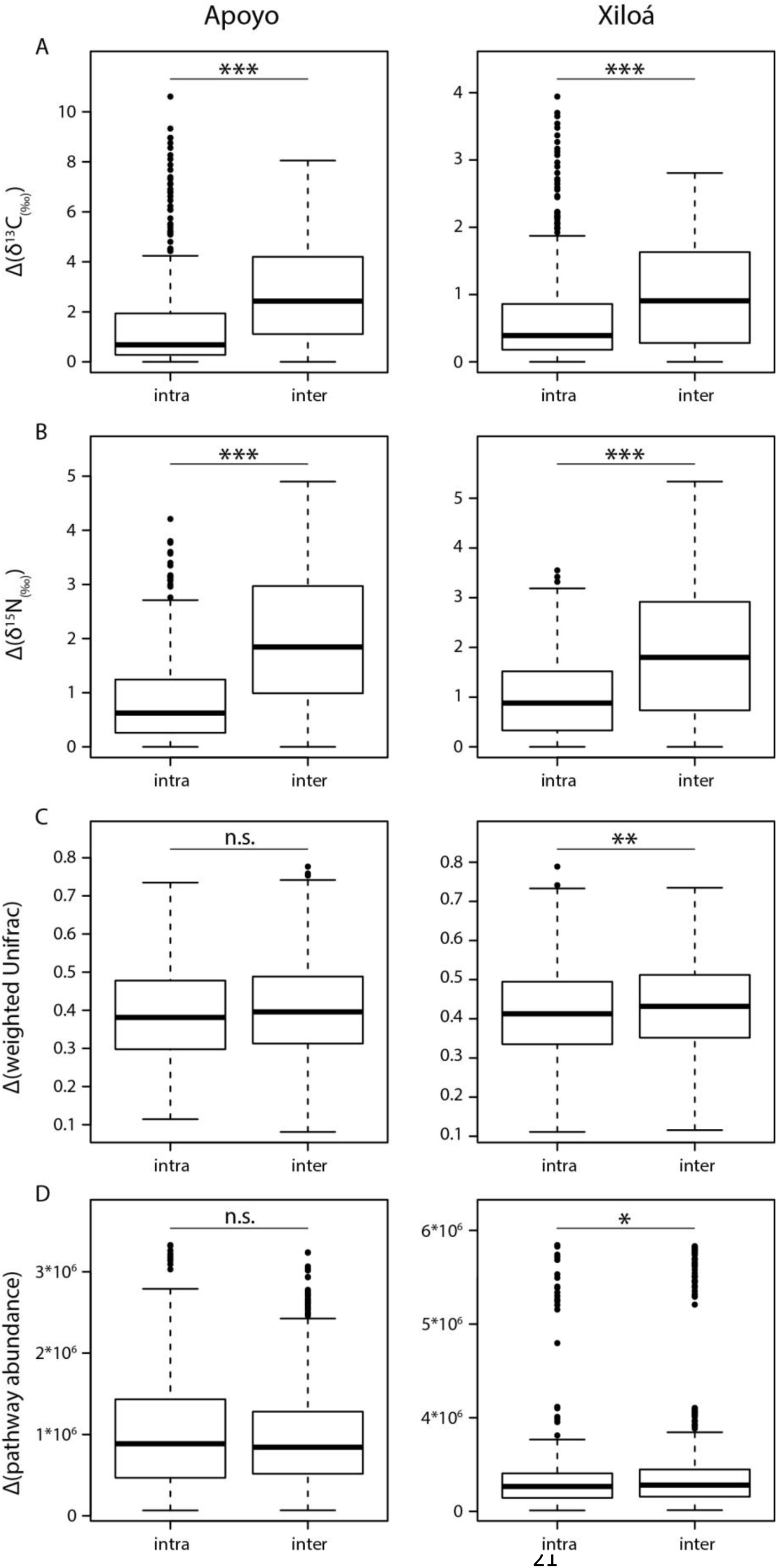
Intra- and interspecific distances in stable isotope values of carbon (A) and nitrogen (B) as well as the gut microbial community (C) and predicted functional bacterial metagenomes (D) among individuals within the two crater lakes. For carbon and nitrogen, interspecific distances are highly significantly larger than intraspecific ones. For the gut microbiota, both taxonomically and functionally, distances are much more similar and significant differences could only be seen in crater lake Xiloá (Wilcoxon rank-sum test, \**P* < 0.05, \*\**P* < 0.01, \*\*\**P* < 0.001).

### Parallelism and non-parallelism of the gut microbiota in crater lake Midas cichlids

Next, we investigated whether the repeated evolution of sympatric species differing in trophic ecology in the crater lakes is associated with parallel changes of the gut microbiota. As δ^15^N values were consistently higher in crater lake Xiloá compared to crater lake Apoyo, we performed z-score normalization of the data to allow comparisons across crater lakes by inferring trophic position (normalized δ^15^N) and littoral carbon usage (normalized δ^13^C) (Fig. S3; 42).

When comparing the adaptive radiations from both crater lakes, bacterial community composition was significantly affected by lake (adonis, *P* = 0.001), but neither by trophic position (*P* = 0.209) nor littoral carbon usage (*P* = 0.857) values. The predicted functional bacterial metagenomes (MetaCyc pathway abundance) was not affected by lake (*P* = 0.191) nor by littoral carbon usage (*P* = 0.521) but was significantly affected by trophic position (*P* = 0.018). Further, we tested which bacterial orders and inferred MetaCyc pathways were affected by trophic ecology in parallel across the two crater lakes (see Methods section for more details). Without correcting for multiple testing, abundances of 21 and nine bacterial orders were affected in parallel by trophic position and littoral carbon usage values, respectively. Likewise, abundances of 15 and two MetaCyc pathways were affected in parallel by trophic position and littoral carbon usage values, respectively. However, this represents only 6.5% (δ^15^N) and 2.8% (δ^13^C) of the 323 bacterial orders and 3.3% (δ^15^N) and 0.4% (δ^13^C) of 459 MetaCyc pathways. None of these bacterial orders or MetaCyc pathways remained significant after correcting for multiple testing (FDR). Taken together, these results indicate that overall gut microbiota differentiation did not occur in parallel with divergence in Midas cichlids’ trophic ecology across the two crater lakes.

## Discussion

Numerous studies on diverse vertebrate species have convincingly demonstrated that the composition of bacterial gut communities is affected by diet (13, 43-45). What remains largely unknown are gut microbiota dynamics during the host’s adaptation to novel food sources, particular during early stages of species divergence. To address this question, we studied trophic ecology and the gut microbiota of repeated Nicaraguan Midas cichlid crater lake radiations, a model system for rapid ecological diversification and speciation (17, 18, 27). We asked whether the parallel evolution of trophic diversification is associated with respective changes of the gut microbiota among sympatric crater lake species (i.e., are the gut microbiotas of ecologically similar species that independently evolved in two crater lakes more similar to each other than they are to their closest relatives of the same lake?). Our results clearly demonstrate that among individuals of the same crater lake differentiation in trophic ecology and the gut microbiota is associated, emphasizing the importance of diet in shaping the gut microbiota. However, we did not find evidence for parallel changes of the gut microbiota across crater lakes, suggesting that diet affected Midas cichlids’ gut microbiota differently in these lakes. Moreover, we found that interspecific variation in trophic ecology (measured as stable isotope ratios of carbon and nitrogen) is significantly higher than intraspecific variation, a pattern that was much less pronounced for the gut microbiota (Fig. 4).

### Gut microbiota differentiation across lakes

Comparing bacterial communities from environmental DNA (eDNA) with those harbored in fish guts clearly revealed that, although bacterial taxa are shared to some extent, the gut microbiota not merely represents the bacterial community of the natural environment but is rather controlled by the host, as has been found for other fishes (14, 15, 46). The dominant bacterial phyla of Midas cichlids’ gut microbiota (Proteobacteria, Firmicutes, Fusobacteria and Bacteroidetes) are also found in many other freshwater fishes (14, 15, 47). Albeit the bacterial species richness of environmental samples strongly differed across lakes (Fig. 3B), the gut microbiota of Midas cichlids, except for *A. citrinellus* from Lake Managua, showed constant levels for this measure. This indicates that the diversity of bacterial species in the gut might be constrained by the host and stabilized at a given level, as predicted by the holobiont theory (48). In *A. citrinellus* from Lake Managua, bacterial species richness was by far the lowest among all populations (Fig. 3B) and was also lower in eDNA from Lake Managua compared to Lake Nicaragua. The city of Managua, Nicaragua’s capital with a population of more than 2 million, is located on the shore of Lake Managua and for decades, domestic and industrial waste water has been disposed into the lake (49). As a result, concentrations of mercury and other toxic substances are extremely high in the lake and are also enriched in fishes (49, 50). Mercury levels have been shown to be correlated with δ^15^N values (51, 52) and Midas cichlids from Lake Managua showed the highest δ^15^N values (Fig. 1), in agreement with the observation that mercury accumulates in these fishes. Albeit speculative at this point, high levels of contamination might have decreased the bacterial species richness of Lake Managua as well as the gut microbiota diversity of Midas cichlids inhabiting this lake.

### Association between trophic ecology and gut microbiota in crater lake adaptive radiations

Diet is widely known to influence the composition of the gut microbiota (13, 44, 45, 56) but most of the previous research mainly focused on model organisms or on distantly related species with long divergence times. However, the dynamics of gut microbiota changes during early stages of ecological diversification and speciation remain largely unknown, but see (14, 15). Midas cichlids represent an excellent model to study such dynamics as species from crater lakes Apoyo and Xiloá repeatedly diverged only very recently and show strong differentiation in trophic ecology (Fig. 1). Therefore, we tested whether gut microbiota differentiation is associated with trophic ecology of crater lake Midas cichlids.

There was a positive correlation between interindividual differentiation of the gut microbiota with carbon isotope signatures in both crater lakes and with nitrogen isotope signatures in crater lake Xiloá (Fig. 4). Accordingly, divergence of trophic ecology and the gut microbiota are to a certain degree associated, suggesting that adaptation to different food sources necessitated changes of the gut microbiota. However, it should be noted that a substantial amount of gut microbiota variation is not explained by trophic divergence. Differences in stable isotope ratios, reflecting the host’s trophic ecology, where considerably higher across species compared to within species (Fig. 5A & B). In contrast, such differences were much less pronounced in crater lake Xiloá and absent in Apoyo for the taxonomic composition of the gut microbiota and the predicted functional bacterial metagenome (Fig. 5C & D). This could mean that occupation of novel trophic niches might be achieved without drastically changing the overall composition of the gut microbiota, both taxonomically and functionally. Rather, subtle changes in some functionally important bacterial taxa might suffice to exploit new food sources and to allow ecological and evolutionary diversification of the hosts. Alternatively, the very recent divergence of crater lake Midas cichlids might impede clear differentiation of the gut microbiota, as discussed in more detail in the following paragraph.

### Parallelism and non-parallelism of the gut microbiota in fishes

Parallel changes of the gut microbiota associated with differentiation in trophic ecology have been reported for older fish species (13), whereas other studies found no evidence for parallelism among more recently diverged populations (14, 15). In the very recent Midas cichlid adaptive radiations from Apoyo and Xiloá that diverged less than 1,700 and 1,300 generations ago, respectively (18), parallel changes in diet led us to expect that a similar pattern could be found in the gut microbiota as well. However, we did not detect evidence for parallel changes of the gut microbiota, except for a significant association of the predicted functional metagenome with trophic position (measured as normalized δ^15^N). Taken together, studies across multiple groups of fishes suggest that parallel changes of the gut microbiota might only be expected on longer time scales (13). These observations can be explained by the fact that during early stages of divergence, species might occupy novel niches but diet, to some extent, still overlaps across young species or ecotypes.

In crater lake Midas cichlids, stable isotope analyses clearly showed that species largely occupy distinct niches (Fig. 1). However, stomach content analyses showed that these species mainly feed on similar food items but their relative proportions differ across species (22). Young crater lake species might be in the process of adapting to specialized ecological niches but currently they are still opportunistic generalists with a varying diet. Thus, short term changes of an individual’s diet are not reflected in the stable isotope signature of muscle tissue as this represents an average of this individual’s diet over a period of approximately three months (57, 58). In contrast, the composition of the gut microbiota is highly variable and changes rapidly with diet (59, 60). Hence, the gut microbiota rather represents a snapshot of an individual’s most recently acquired food items, generating high levels of intraspecific variation. This could explain why intra- and interspecific variation of the gut microbiota is much more equal compared to stable isotope data (Fig. 5). Accordingly, high levels of dietary intraspecific variation might mask interspecific differences in trophic ecology, thereby blurring any signal of gut microbiota parallelism in recently diverged ecotypes or species. This is what we can also see in other fishes like whitefish and guppies, where the main change between ecotypes is in the relative proportion of food items (14, 15). Only after species sufficiently diverged to become trophic specialists that do not overlap in food items (e.g., carnivorous and herbivorous cichlids from Lake Tanganyika), one would expect persistent and parallel patterns of gut microbiota divergence, as seen in African cichlids (13). Thus, taking into account ecological characteristics (e.g., extent of trophic divergence) as well as the evolutionary history of host species will aid in predicting when to expect parallel changes of the gut microbiota. This, in turn, will improve our understanding of gut microbiota dynamics during early stages of ecological diversification.

To conclude, in this study we illustrate that (i) the gut microbiota of Midas cichlids from distinct environments is clearly differentiated, (ii) within crater lakes, divergence of the gut microbiota is associated with trophic ecology but such changes did not occur in parallel across crater lakes and (iii) differentiation in trophic ecology is not reflected by large rearrangements of gut bacterial communities in these recently diverged species.

## Acknowledgements

We thank the Ministry of Natural Resources (MARENA) in Nicaragua for collection and exportation permits. Further, we would like to thank L. Paíz-Medina and R. Rayo for help in the field and W. Kornberger for aiding in stable isotope analyses. This work was supported by the European Research Council through ERC-advanced (grant number 293700-GenAdap to A.M.) and the University of Konstanz.

## Author contributions

A.H., J.T.D. and A.M. developed the project; A.H., J.T.D., G.M.S. and A.M. collected samples in the field. A.H. extracted DNA, prepared the libraries and analyzed the sequencing data. A.H. and S.R. prepared samples for stable isotope analyses. Stable isotope data was collected by E.Y. and analyzed by A.H., J.T.D., S.R. and E.Y.; A.H. wrote the manuscript with input from all of the authors.

