## Supplementary Figures 1-3 for "Parallel and non-parallel changes of the gut microbiota during trophic diversification in repeated young adaptive radiations of sympatric cichlid fish"

**Supplementary Material**

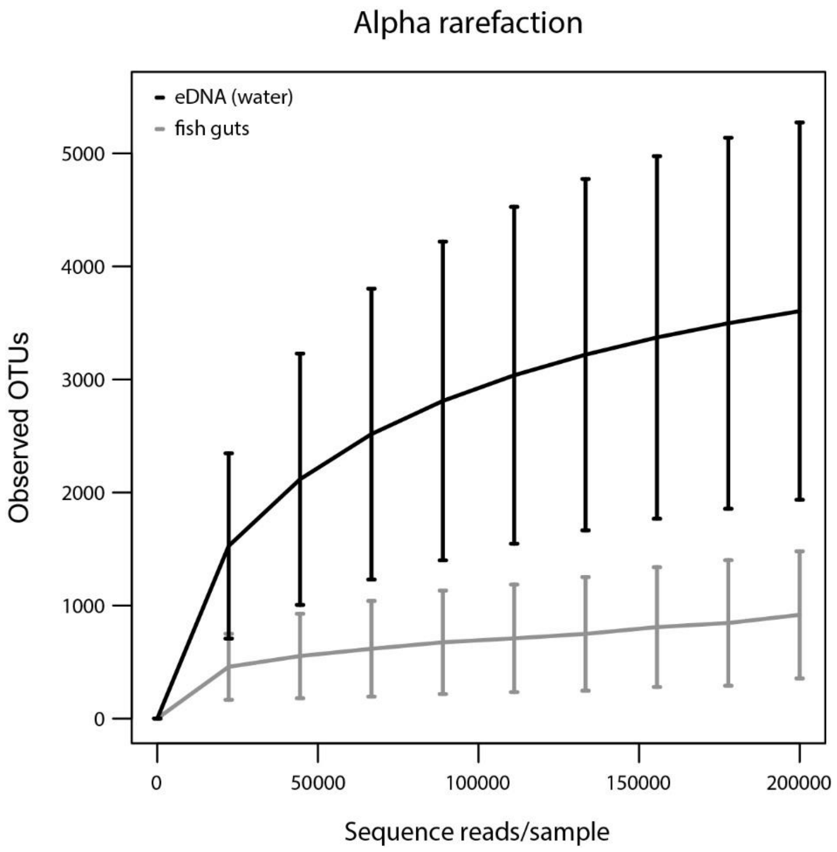


Supplementary Figure 1: Alpha diversity estimates at different rarefaction depths for eDNA (black) and fish (grey) samples. The investigated sequencing depths range from 11 to 200,000 reads. At a sampling depth of 20,000 reads, a large proportion of the microbial diversity in fish guts is captured.

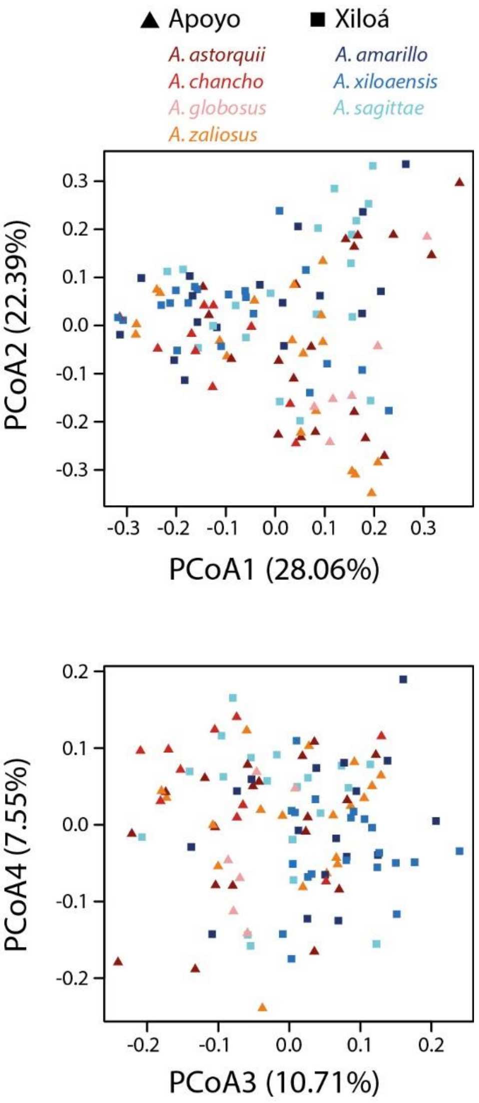


Supplementary Figure 2: Principal coordinate analysis of gut microbiota from crater lake Midas cichlids measured as weighted UniFrac. We could not detect any apparent clustering by lake or between benthic and limnetic species along PCoAs 1 to 4.

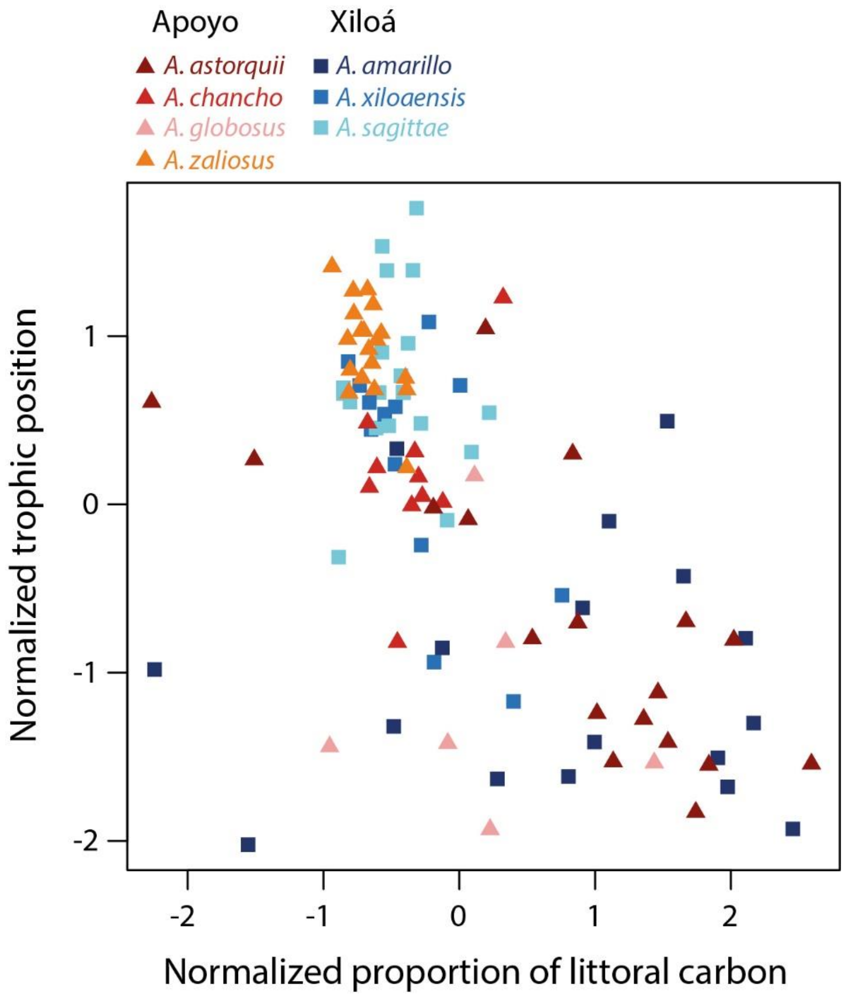


Supplementary Figure 3: Trophic position and proportion of littoral carbon of crater lake Midas cichlids were inferred by performing a z-normalization of nitrogen and carbon stable isotope values.
